# Antiretroviral therapy is a major cause of tuberculosis decline in southern and eastern Africa

**DOI:** 10.1101/477158

**Authors:** Christopher Dye, Brian G. Williams

## Abstract

The incidence of tuberculosis (TB) in southern and eastern Africa was driven sharply upwards during 1980s and 1990s by coinfection with *Mycobacterium tuberculosis* and the human immunodeficiency virus (HIV). Although drug treatments for TB infection (isoniazid preventive therapy) and disease (combinations of TB drugs) can reduce TB incidence if implemented effectively, we find that antiretroviral therapy (ART) given to people with HIV infection was strongly and systematically associated with the accelerated decline of TB in 12 of the worst affected African countries between 2003 and 2016. Inferring that ART was a significant cause of TB decline, ART prevented approximately 1.88 ± 0.23 million HIV-positive TB cases, or 15.7 ± 1.9 percent of the total number expected. There is no evidence that drug treatment of TB infection (IPT) or disease (combination chemotherapy) played more than a minor role in accelerating TB decline after 2003. In these 12 countries, over the period 2003–16, ART made a major contribution towards achieving international targets for the reduction of TB incidence.

**Significance:** Tuberculosis (TB) is still the leading cause of death from a single infectious agent. To cut the TB incidence rate by 80% between 2015 and 2030, in line with the WHO End TB Strategy, demands a five-fold increase in the rate of decline worldwide, from 2% to 10%/year. We find that the reduction in TB incidence rate in 12 African countries, at up to 8%/year, is due mainly to the expanded provision of antiretroviral therapy (ART) to people living with HIV, rather than to improvements in the treatment of TB infection and disease. ART should remain central to TB control where rates of TB-HIV coinfection are high, but new efforts are needed to maximize the direct benefits of treating TB infection and disease.

---

Among people infected with *Mycobacterium tuberculosis*, co-infection with the human immunodeficiency virus (HIV) carries a 10- to 15-fold risk of developing tuberculosis disease (TB) (1–3). As HIV infection spread through the populations of southern and eastern Africa from the 1980s onwards, the worst affected countries, including Botswana, Lesotho and Swaziland, have reported national adult (15–49 years) HIV prevalence rates over 20% and exceptionally high TB incidence rates above 500/100,000 population/year (Table 1, columns 3 and 5). Across the whole of Africa in 2016, there were an estimated 764,000 new cases of TB among people living with HIV (PLHIV) leading to 320,000 deaths (4).

**Table 1.** TB and HIV epidemics, and interventions to reduce TB incidence, in 12 countries of southern and eastern Africa, 2003–16.

| Location |  | TB and HIV burden and trends |  |  |  |  |  |  | Interventions |  |  |  |  |  |
| --- | --- | --- | --- | --- | --- | --- | --- | --- | --- | --- | --- | --- | --- | --- |
| Country | Code | HIV prevalence (15-49 yr, max *H*) | HIV prevalence in TB (% max) | TB incidence (per 100,000 population, max) | HIV+ TB incidence (per 100,000 population, max) | Change HIV+ TB incidence 2003-16 (%/year) | Change HIV- TB incidence 2003-16 (%/year) | Change total TB incidence 2003-16 (%/year) | ART coverage 2003-16 (%PLHIV, *A*) | Average TB detected and successfully treated 2003-16 (%) | Change TB detected and successfully treated 2003-16 (%) | IPT coverage 2003-16 (%PLHIV) | HIV+ TB cases prevented (000s) | HIV+ TB cases prevented (%) |
| 1 | 2 | 3 | 4 | 5 | 6 | 7 | 8 | 9 | 10 | 11 | 12 | 13 | 14 | 15 |
| Botswana | BWA | 0.24 | 70 | 816 | 571 | -8.1 | -4.0 | -6.9 | 34 | 51 | 12.5 | 0.51 | 48 | 30 |
| Kenya | KEN | 0.08 | 51 | 397 | 331 | -7.6 | -1.2 | -3.8 | 19 | 38 | 6.3 | 0.56 | 392 | 21 |
| Lesotho | LSO | 0.24 | 77 | 1280 | 979 | -4.1 | -2.1 | -3.7 | 16 | 33 | 10.4 | 0.11 | 75 | 23 |
| Malawi | MWI | 0.13 | 71 | 397 | 283 | -7.5 | -3.1 | -5.9 | 20 | 51 | -9.0 | 3.09 | 111 | 19 |
| Namibia | NAM | 0.14 | 59 | 935 | 552 | -9.0 | -2.5 | -5.7 | 26 | 61 | 30.5 | 0.44 | 55 | 31 |
| Rwanda | RWA | 0.04 | 45 | 102 | 46 | -5.2 | -3.1 | -4.7 | 31 | 62 | 14.8 | 0.01 | 19 | 30 |
| South Africa | ZAF | 0.17 | 65 | 977 | 607 | -0.5 | -0.1 | -0.4 | 14 | 47 | 28.2 | 4.21 | 543 | 11 |
| Swaziland | SWZ | 0.27 | 84 | 1280 | 973 | -6.8 | -2.2 | -5.8 | 22 | 39 | 91.9 | 0.17 | 54 | 27 |
| Tanzania | TZA | 0.07 | 47 | 510 | 243 | -7.1 | -2.7 | -4.4 | 15 | 28 | 16.9 | 0.18 | 195 | 13 |
| Uganda | UGA | 0.08 | 59 | 248 | 146 | -4.3 | 1.0 | -1.7 | 18 | 44 | -0.4 | 0.001 | 72 | 11 |
| Zambia | ZMB | 0.14 | 70 | 662 | 445 | -5.3 | -2.7 | -4.3 | 23 | 56 | -7.5 | 0.02 | 165 | 20 |
| Zimbabwe | ZWE | 0.19 | 78 | 617 | 471 | -8.7 | -6.0 | -7.8 | 19 | 49 | 21.1 | 2.93 | 149 | 18 |

To reduce the incidence of TB in populations with high rates of *M. tuberculosis* and HIV coinfection, the World Health Organization (WHO) and the Joint United Nations Programme on HIV/AIDS (UNAIDS) recommend primarily drug treatments to prevent TB disease (isoniazid preventive therapy, IPT, acting on *M. tuberculosis* infection, and antiretroviral therapy, ART, acting on HIV infection) and to prevent the further transmission of infection from HIV-positive and HIV-negative patients with TB disease (combinations of TB drugs) (4, 5). The efficacy of these treatments has been demonstrated in experimental and observational studies (6–11) and they have the potential markedly to reduce incidence nationally if high rates of coverage can be achieved in target populations (1, 2, 12–14).

In view of these empirical studies and forecasts, the purpose of the present retrospective analysis is to investigate, drawing on data collected annually since 1990, which of these recommended methods have in practice had the greatest impact on TB incidence in 12 African countries that carry high burdens of TB linked to HIV infection (SI Appendix, Fig. S1).

Fig. 1 illustrates for two countries, Botswana and South Africa, the time course of TB and HIV epidemics and their control measures between 1990 and 2016 (epidemic trajectories for all 12 countries are shown in the SI Appendix, Fig S2). The numbers of TB cases reported each year increased following the rise of HIV incidence and prevalence. The subsequent downturn in TB case incidence in both countries appeared to be associated with improvements in the proportion of TB cases successfully treated (i.e. proportion of cases detected × proportion successfully treated), but more obviously with the expanded coverage of ART. The scale-up of ART coverage from 2003 onwards was more rapid in Botswana than in South Africa (Fig. 1), and the decline in TB incidence occurred earlier and more quickly. In these two countries (and all others investigated here) the proportional decline, and the absolute magnitude of the decline, was greater for HIV-positive than for HIV-negative TB cases (Table 1, columns 7 and 8). The provision of IPT, compared with ART for PLHIV and the treatment of TB patients, was very low in both countries (Fig. 1).

**Fig. 1.**
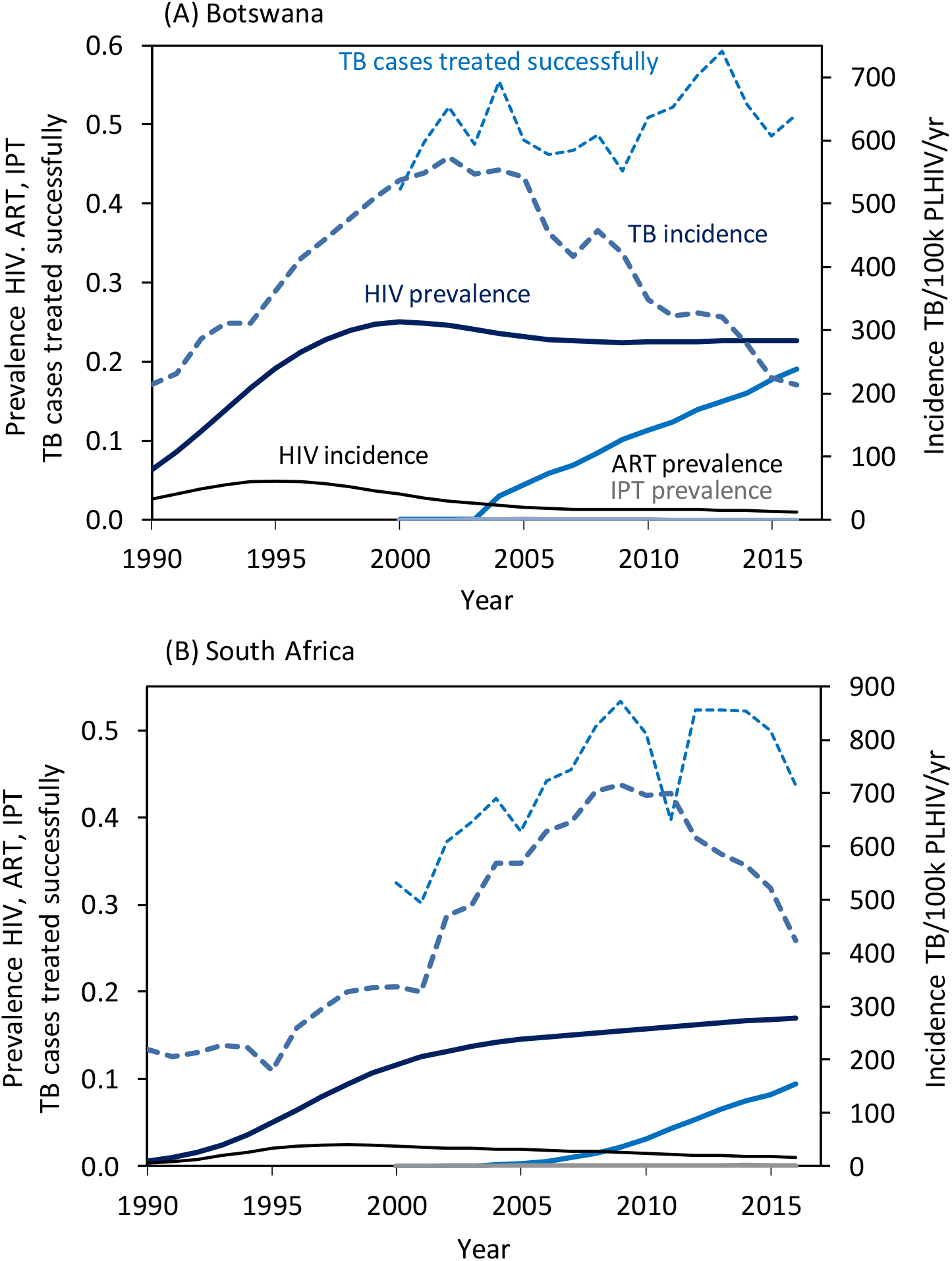
Time trends of TB and HIV epidemics in southern and eastern Africa, 1990–2016, illustrated by (A) Botswana and (B) South Africa. Continuous lines (*y*-axis left) show trends in estimated HIV incidence and prevalence in adults (15–49 years), and ART and IPT prevalence (coverage in the whole population). Broken lines show the reported TB incidence (y-axis right) and the proportion of cases detected and successfully treated, i.e. the product of proportions of cases detected and successfully treated (y-axis left). Data are from WHO and UNAIDS (4, 23).

To examine the effect of control measures systematically across all 12 countries, we used simple mathematical models and routinely collected data to investigate the effects of ART and IPT (Fig. 2) in preventing TB disease in PLHIV, and to investigate TB case detection and treatment (Fig. 3) as a means of interrupting *M. tuberculosis* transmission and consequently reducing disease incidence (Materials and Methods). There appear to be no other competing explanations for the accelerated rate of decline in TB in these 12 countries from the early 2000s onwards.

**Fig. 2.**
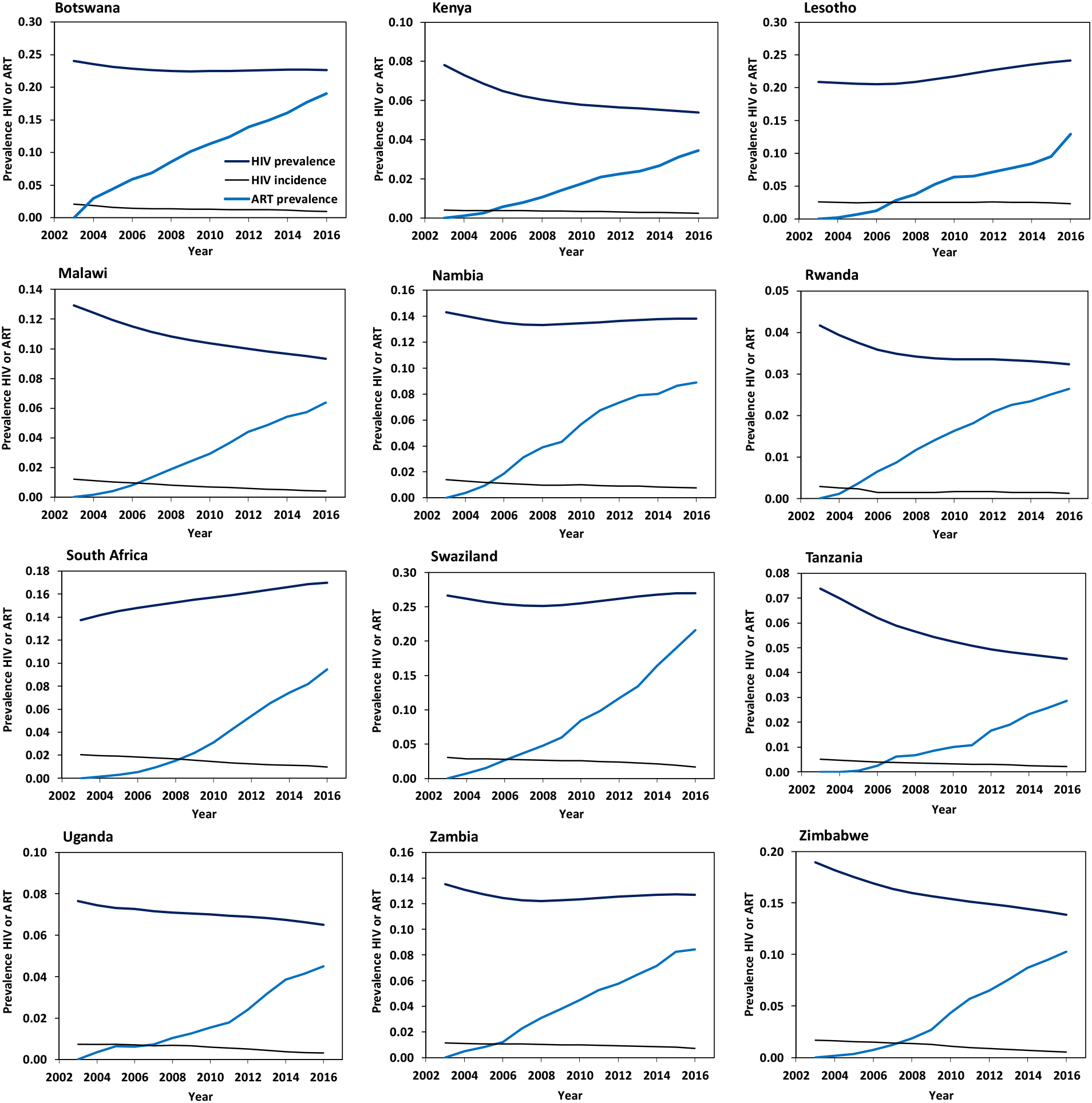
Time trends in HIV incidence, prevalence and ART coverage in the populations of 12 countries in southern and eastern Africa, 2003–16. The ratio of ART/HIV prevalence is the coverage of ART in PLHIV.

**Fig. 3.**
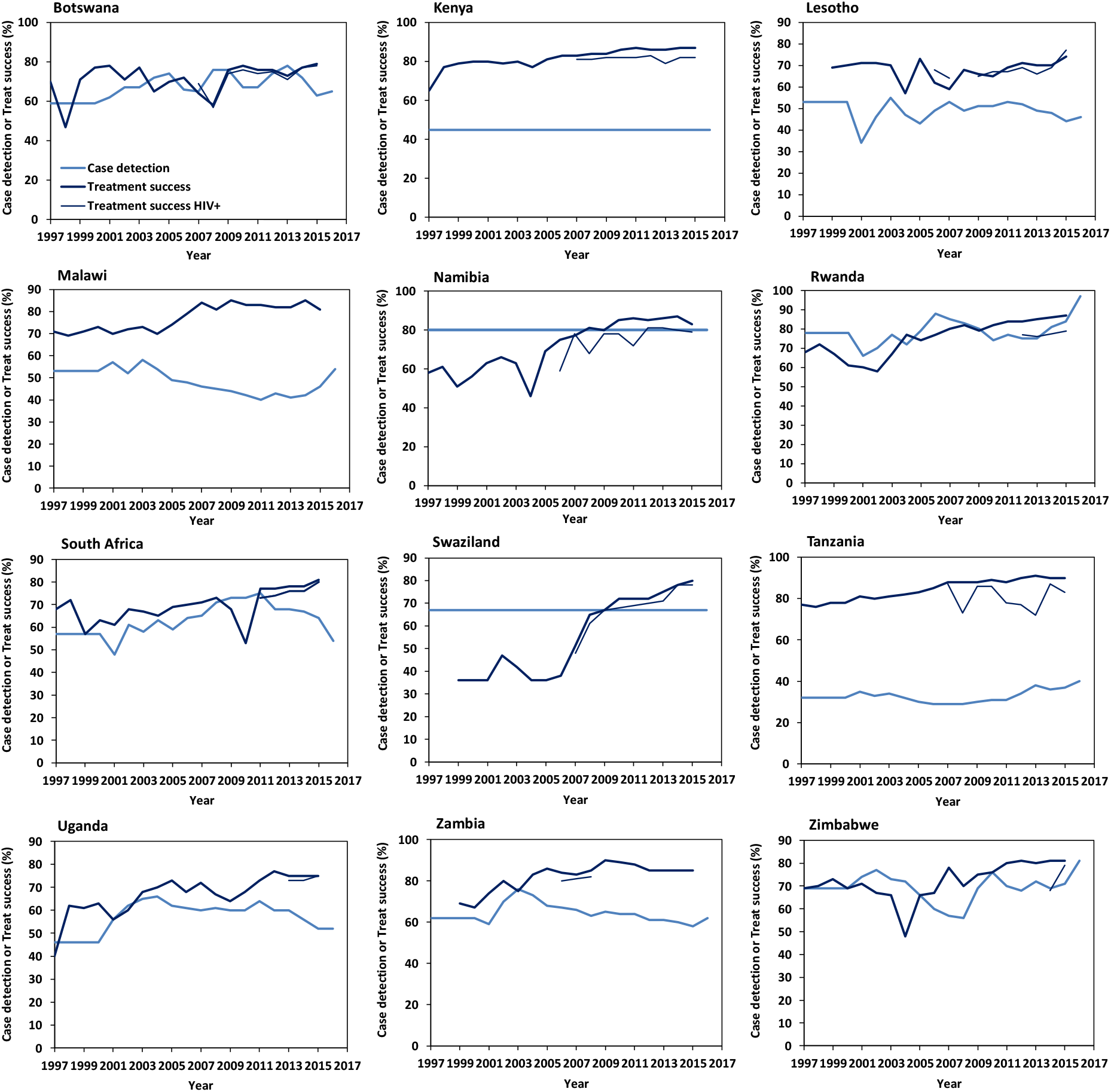
Time trends in TB case detection and treatment success for 12 countries in southern and eastern Africa. Data are shown for the period 1997–2016 although the analysis focuses on the period 2003–16. Data are the estimated percentage of cases detected and the measured proportion of cases successfully treated for all forms of TB, and treatment success for HIV-positive TB cases where data are available.

## Results

ART coverage increased in all countries between 2003 and 2016 (Fig. 2), while TB incidence was in decline (Figs. 1 and 4; SI Appendix, Fig. S2) so the measured changes in both are inevitably correlated (SI Appendix, Table S1). Considering each country separately, these correlations do not prove that TB decline was caused by ART because the associations could be confounded by other unmeasured determinants that also change with time. However, the evidence for a causal role of ART becomes compelling when the results are compiled for all 12 countries: modelling the change in HIV-positive TB incidence with and without ART (Materials and Methods, Fig. 4) reveals that the estimated fraction of HIV-positive TB cases prevented by ART was strongly and systematically associated with the provision of ART (Fig. 5, *r*^2^ = 0.68, *p* = 0.0009).

**Fig. 4.**
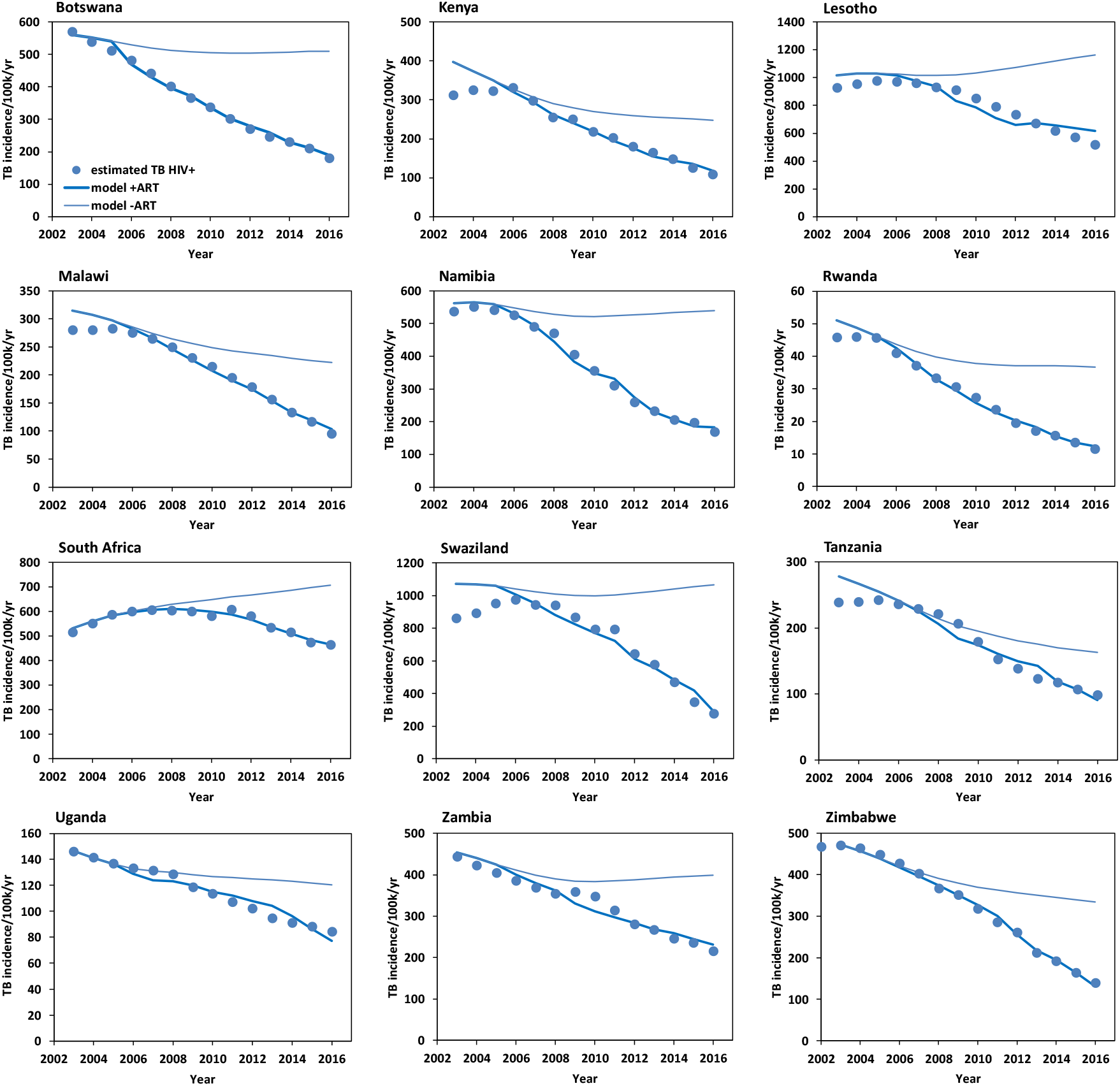
Time trends in HIV-positive TB incidence. Data points are the estimated incidence of TB cases/100,000 population/year; solid lines represent the fit of the model under the hypothesis that ART prevents (for regression coefficients β1 > 0, lower lines) or does not prevent TB (counterfactual β = 0, upper lines). The number of cases prevented in each country, 2003–16, is calculated from the area between upper and lower lines.

ART coverage was highest on average, and the greatest fractions of HIV-positive TB cases were prevented, in Botswana, Namibia and Rwanda (Table 1, columns 10 and 15). South Africa prevented the greatest number of HIV-positive TB cases (543,000) between 2003 and 2016, being the country with the largest number of PLHIV (7.1 million in 2016). But South Africa prevented the smallest fraction of cases (11%) because ART coverage was relatively low on average (22%, Table 1).

Based on the fit of the model to data for each of the 12 countries (Fig. 4; SI Appendix, Fig S5), a total of 1.88 ± 0.23 million HIV-positive TB cases were prevented by ART over the period 2003 to 2016, or 15.7 ± 1.9 percent of the total number expected. An alternative way to calculate the number of TB cases prevented is from the systematic, region-wide association with ART coverage (Fig. 5): summing the predictions for each country gives a similar total of 1.92 million HIV-positive TB cases prevented between 2003 and 2016, or 17.3 percent of the total expected.

**Fig. 5.**
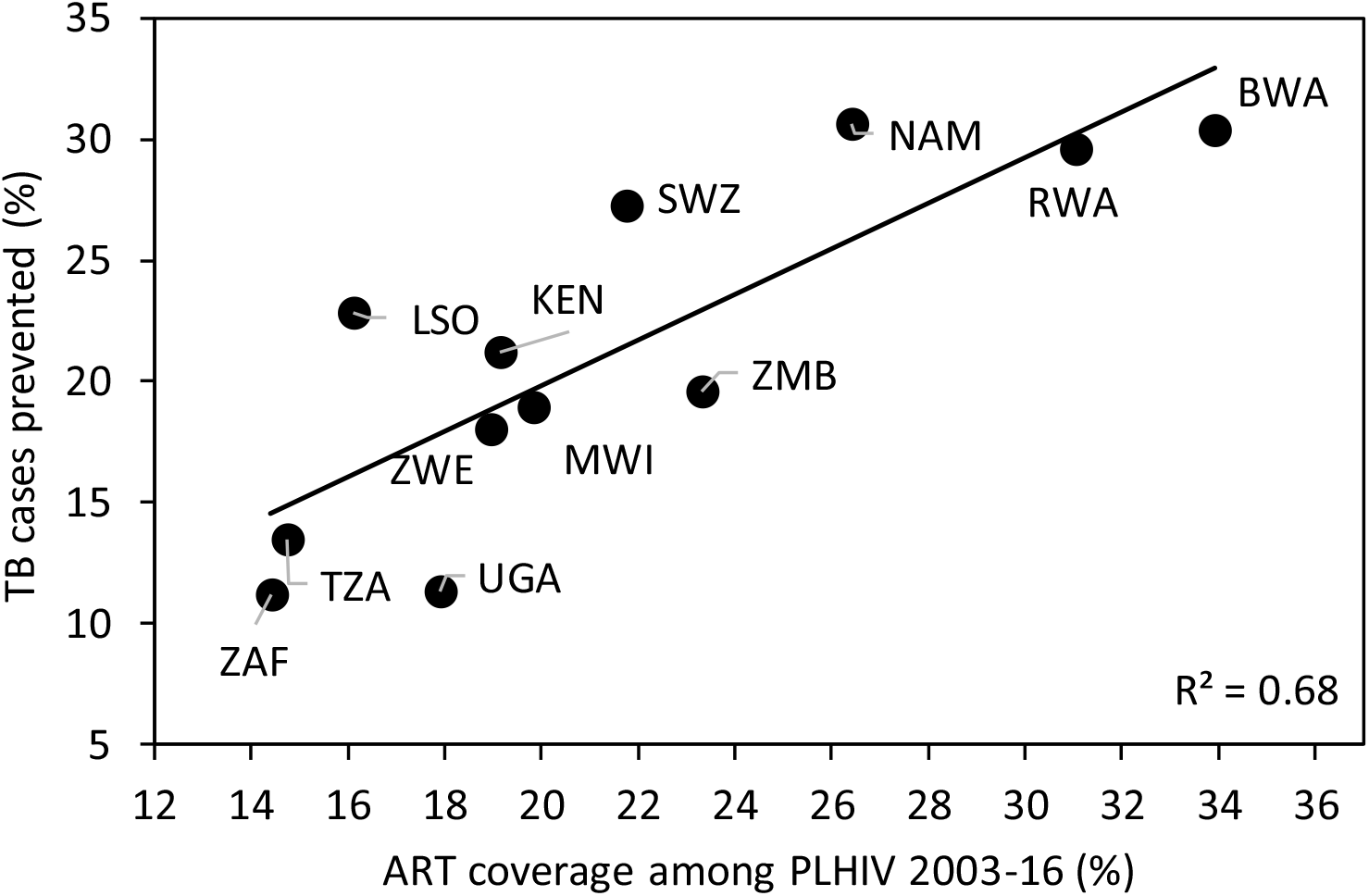
The association between ART coverage and the percentage of HIV-positive TB cases prevented in 12 countries of southern and eastern Africa, 2003–16, assuming a causal role for ART. ART coverage explains 68% of the variation in the estimated percentage of HIV-positive TB cases prevented (*r*^2^ = 0.68, p = 0.0009). Full country names are listed in Table 1.

Although the primary effect of ART is to stop progression from *M. tuberculosis* infection to TB disease in PLHIV, the consequent reduction in incidence might also reduce onward transmission. In this context, we observe that the incidence of HIV-negative TB was falling in all countries alongside HIV-positive TB, albeit more slowly (Table 1, columns 7 and 8) and, comparing countries, the rates of decline are correlated (*r*^2^ = 0.48, *p* = 0.02). The possibility that ART reduces *M. tuberculosis* transmission as well as disease progression remains to be proven; if ART does help to interrupt transmission to HIV-positive and HIV-negative individuls, our estimate of about 1.9 million TB cases prevented would be conservative.

Considering the possible role of other interventions, there is little evidence that the treatment of TB directly, either prophylactic drug treatment of *M. tuberculosis* infection (IPT) or the treatment of TB disease with combinations of drugs, contributed substantially to the accelerated decline in TB incidence between 2003 and 2016.

First, the detection of infectious, pulmonary TB cases early in the course of illness, followed by successful drug treatment, should interrupt the transmission of *M. tuberculosis* from both HIV-positive and HIV-negative cases, thereby reducing TB incidence in the rest of the population (3, 12, 14). Comparing countries, the rate of decline in TB incidence is expected to be faster where case detection and treatment success are higher on average. The national proportions of all TB cases detected and successfully treated were generally low, and varied between 28% and 62% (Table 1, column 11), but there was no discernible association across the 12 countries investigated here (Table 1, columns 9 and 11; *r*^2^ = 0.06, *p* > 0.05).

If case detection and treatment success improve with time in any individual country, we expect to see an accelerated decline in TB incidence as the interruption of transmission lowers the TB case reproduction number (Materials and Methods). TB case detection and treatment did improve somewhat in the majority of countries between 2003 and 2016 as TB incidence declined, though not in Malawi, Uganda and Zambia (Table 1, column 12; Fig. 3). However, within each country, there were either weak associations (South Africa, Swaziland, Zimbabwe) or no significant associations (the other nine countries) between TB detection and treatment success and the rate of decline in TB incidence (SI Appendix, Table S1). In short, however we examine the measured rates of TB detection and treatment success, the data do not explain the systematic, accelerated reduction in TB incidence observed in all 12 countries from around 2003 onwards (Fig. 4).

Second, IPT used continuously with ART generally gives coinfected adults additional protection from TB, augmenting that provided by ART alone (9–11, 15). In the 12 countries investigated here, the coverage of IPT among PLHIV over the period 2003–16 was generally low (Table 1). South Africa started the largest number of people on IPT in 2016, 385,932 or 51% of those newly enrolled in HIV/AIDS care in that year. But only 4.2% of PLHIV, on average, received IPT between 2003 and 2016, compared with 22% who started ART. Behind South Africa, 3.1% of PLHIV received IPT in Malawi and 2.9% in Zimbabwe. These three countries together reported 91% of all PLHIV who started IPT in the 12 countries (Table 1). Thus IPT may have protected thousands of coinfected people from developing TB in Malawi, South Africa and Zimbabwe, but the effect on TB incidence at population level would have been relatively small in these three countries, and negligible in the other nine.

## Discussion

Our analysis suggests that the roll-out of ART has played a major part in accelerating the rate of TB decline in southern and eastern Africa since 2003; the greater the coverage, the greater the impact. In contrast, there is no evidence that drug treatment of TB infection (IPT) or disease (combination chemotherapy) played more than a minor role in accelerating the rate of TB decline.

We found large variations among countries in the estimated fraction of TB cases prevented by ART, linked to differences in ART coverage. Both the immediate and deeper reasons for these differences in coverage need to be understood because they are likely to influence the future success of HIV and TB control programmes. Botswana prevented the largest fraction of TB cases between 2003 and 2016 by achieving the highest coverage of ART. Since independence in 1966, Botswana has invested in public services to tackle ill health and poverty (16) and, in that broader context, was the first African country to provide universal free ART (17). South Africa prevented the smallest fraction of TB cases among the 12 countries investigated here. Following controversy over the link between HIV and AIDS, South Africa was slower to expand treatment for PLHIV, but now has the largest number of people on ART and IPT worldwide. More recently, South Africa was the first country in the region to approve pre-exposure prophylaxis (PrEP), in which ART protects HIV-negative people from acquiring infection. Consistent with these recent developments, the rate of TB decline among PLHIV increased to 4.6%/year between 2010 and 2016 (cf the average of 0.5%/year over the period 2003–16, Table 1, column 7).

Concerning the detection and treatment of TB cases, the mainstay of TB control worldwide, the most important recent trend in Africa has been a sharp growth in the number of TB cases tested for HIV infection. Across Africa in 2016, 82% of reported TB cases had a documented HIV test result, up from just 2% in 2004 (4). Having been tested for HIV, TB cases become eligible for ART, increasing the chance of successful treatment and reducing the risk of death. But far less progress has been made in finding and treating TB among PLHIV, and among HIV-negative people living in settings with high rates of coinfection. And yet both are imperative for TB control, and ultimately for TB elimination (3). To find infectious TB cases earlier among people with and without HIV infection requires a greater awareness of TB and HIV among people at risk, and a more active approach to TB diagnosis at home, in the community and in health facilities (18–21).

With regard to TB preventive therapy, while millions of PLHIV now receive ART continuously in southern and eastern Africa, a relatively small number also receive IPT (4, 15). In this context, South Africa’s success in starting more than 350,000 patients on IPT in 2016, over half of those enrolled in HIV/AIDS care in that year, will add wisdom on how to identify candidates for IPT (recognizing that efficacy is greater for those who are tuberculin skin test positive, and that patients with TB disease must be excluded), on how to maintain adherence to prolonged prophylaxis (typically, daily treatment for 6 months), and on how to manage the liver toxicity that affects a small fraction of patients on isoniazid (22).

## Conclusion

TB is still the largest cause of death from a single infectious agent worldwide (4) and TB incidence is falling far too slowly to reach international targets. The WHO End TB Strategy aims to cut the TB incidence rate worldwide by 80% between 2015 and 2030, which requires a five-fold increase in the rate of decline, from 2%/year at present to at least 10%/year. The slow pace of technological development in TB control — including diagnostics, drugs and vaccines (4) — is a strong argument for making best use of all the tools that are currently available. The best strategy for doing so will vary from one setting to another. But for populations suffering high rates of TB-HIV coinfection, including the 12 countries in this study, ART is evidently a leading intervention. But the scale-up of ART alone will not be enough to reach international targets; besides the search for new interventions and technologies, additional efforts are needed to maximize the well-established, direct benefits of treating TB infection and disease.

## Materials and Methods

### Data sources

The TB and HIV data used in this analysis are from the World Health Organization (WHO) (4) and the Joint United Nations Programme on HIV/AIDS (UNAIDS) (23), and can be downloaded from www.who.int/tb/data/en/ and www.unaids.org/en/resources/documents/2017/2017_data_book. Population data are from esa.un.org/unpd/wpp/. These data can also be obtained from data.worldbank.org.

### Countries included in the analysis

We chose 12 countries of southern and eastern Africa that have high TB-HIV coinfection rates per capita and sufficient data for this investigation (SI Appendix, Fig. S1). The time course of TB and HIV epidemics in these countries follows the general pattern described for Botswana and South Africa in the main text (SI Appendix, Fig S2). We excluded one large contiguous country, Mozambique, because the time trend in TB case incidence is unclear from the series of annual case notifications: the number of cases reported was increasing continuously between 2002 and 2016, apparently because the proportion of incident cases detected was also increasing over that period, and not because of a true rise in incidence (SI Appendix, Fig S3).

### Modelling interventions to reduce TB incidence

The two principal interventions to reduce TB incidence — those with greatest population coverage and longest history of use — are ART to stop progression from infection to disease, and combination chemotherapy to prevent onward transmission from infectious TB cases (the coverage of isoniazid preventive therapy has so far been much lower). Because these interventions to stop progression and transmission have different modes of action, we have investigated their effects using two separate univariate models, rather than combining them in one multivariate model.

(a) Modelling the effect of ART on TB incidence. The annual incidence rate of TB among PLHIV in year *τ* can be estimated from *T_t_ = αH_t−τ_*(1 − *βA_t−τ_*). In this model *α* is the per capita rate at which TB cases arise in the untreated population of PLHIV (*H*) after a delay of *τ* years (i.e. the incidence of TB disease), which depends on the prevalence and incubation period of *M. tuberculosis* in HIV-positive individuals and on the average age of HIV infection in PLHIV. In the absence of drug treatment of either *M. tuberculosis* or HIV infection, *α* would change little over the period 2003–16, so we take *α* to be constant over the time span of this study. *A* is the fraction of PLHIV receiving ART, which takes effect after ***τ*** years, and coefficient *β* scales the effect of ART under the hypothesis that ART prevents TB. This model, and this analysis, do not explore the separate effect of ART on HIV, which is to interrupt viral transmission and to increase population prevalence by greatly improving survival and life expectancy (24–26).

For each of the 12 countries, *T* is obtained from routine surveillance data (annual TB case notifications), and from estimates of the proportions of cases detected and infected with HIV (4), and *H* and *A* from national estimates derived from national and subnational survey data (23). Rewriting the model as *T_t_/H_t−τ_ = α − αβA_t−τ_*, parameters *α* and *β* are estimated from a linear regression of *T_t_/H_t−τ_* on *A*, with intercept *α* and gradient *−αβ*. The time delay, *τ*, is set at 2 years, based on the fit of the regression model to data, and on the recovery rate of CD4 cell count after patients are started on ART (27) (SI Appendix). The number of HIV-positive TB cases prevented is obtained by comparing the numbers expected (modelled) under the hypothesis that ART prevents (*β* > 0, estimated) or does not prevent (*β* = 0, counterfactual) TB.

The length of the time delay, *τ*, depends on the time taken for ART (*A_t−τ_*) to become protective, thereby stopping coinfected people from progressing to TB disease. Fitting the above model to data by linear regression shows that there is a delay between the introduction of ART and its effect in reducing TB incidence — TB incidence continues briefly to rise before falling. Setting *τ* = 2 years eliminates the effect of this delay, as illustrated for Kenya and Swaziland in the SI Appendix (Fig. S4). The fit of the model for all countries with *τ* = 2 years is in Fig. S5. A delay of 2 years is also justified by the study of Gras *et al* (27) who found that, once HIV-positive people start ART, their CD4 cell counts increase at a rate that is independent of the initial CD4 count (Fig. S6A), with a characteristic recovery time of 1.65 years (≈ 2 years) (SI Appendix, Fig. S6B). There is also a delay from the time at which people become coinfected and the onset of TB, which depends on the incubation period of *M. tuberculosis* in HIV-positive individuals and on the average age of HIV infection in PLHIV. When a person infected with *M. tuberculosis* also acquires HIV (coinfection), TB disease will develop after an incubation period of 6–8 years (1, 3). In African countries now, PLHIV are carrying HIV infections of mixed ages — some acquired recently and some acquired many years ago — so people who are coinfected now are part of the way through the incubation period. For convenience, we used the same delay of *τ* = 2 years to relate *T_t_* to *H_t−τ_* as we used to relate *T_t_* to *A_t−τ_* (above).

(b) Modelling the effect of TB case detection and treatment on TB incidence. We evaluated the potential effect of TB case detection and successful treatment on TB incidence, via reduced transmission, using the model *T_t_ = rT_t−τ_*(1 − *δD_t−τ_*), where *T_t_* and *T_t−τ_* represent case incidence in the whole TB and human populations. Coefficient *r* is the TB case reproduction number over the period *τ* in the absence of drug treatment. *D_t−τ_* is the proportion of all TB cases detected and successfully treated and *δ* scales the effect of *D_t−τ_* in reducing *M. tuberculosis* transmission. Rewriting the model as *T_t_/T_t−τ_ = r − rδD_t−τ_*, parameters *r* and *δ* are estimated from a linear regression of *T_t_/T_t−τ_* on D, with intercept *r* and gradient -*rδ*. The regression analysis is a test of whether an increase in TB detection and treatment accelerates the decline in TB incidence by lowering the case reproduction number. *D* is calculated from estimates of the proportion of all TB cases detected in any given year multiplied by the proportion treated successfully, which is reported by WHO in the following year (4). The time delay, *τ*, is set at 2 years, as for the ART model, but the conclusions of the analysis are not sensitive to the length of the delay.

## Supporting information

## Acknowledgments

Funding: none. Author contributions: CD conceived the study, carried out preliminary analysis and drafted the paper. Both authors compiled the data, carried out the full analysis and interpretation, and finalised the text. Competing interests: the authors declare no competing interests.

