## Supplementary material for "Antiretroviral therapy is a major cause of tuberculosis decline in southern and eastern Africa"

The SI presents additional data on HIV and TB epidemics, and additional Figures to support Materials and Methods in the main text.

**Countries included in the analysis**

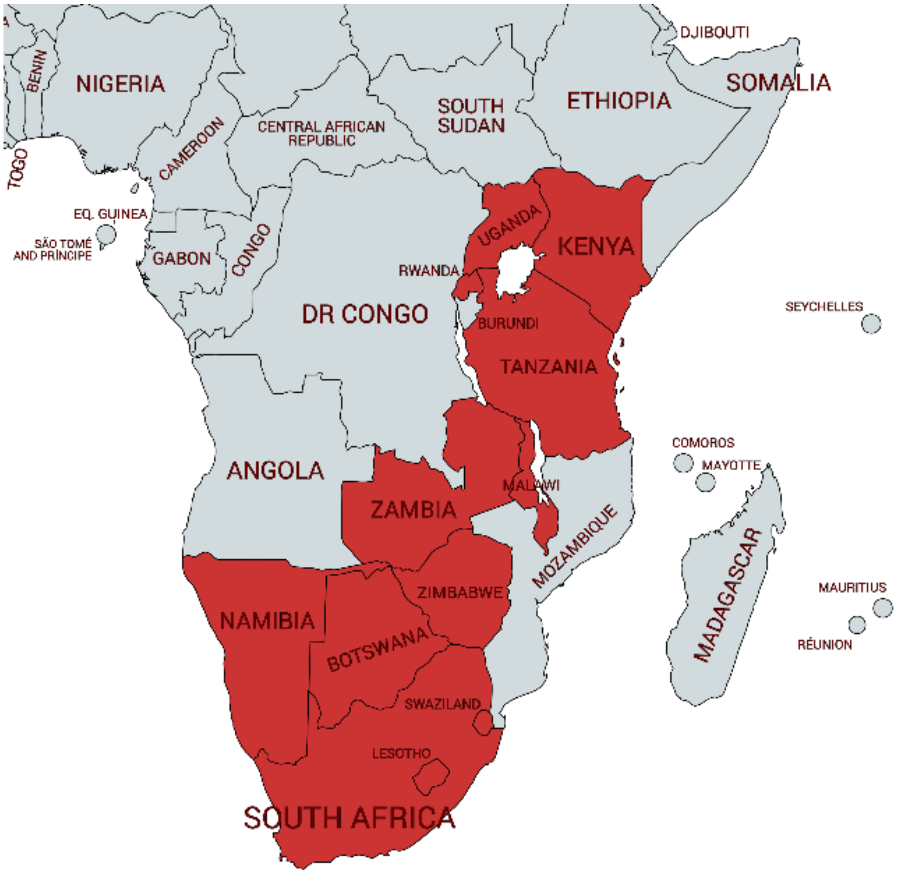


Fig. S1. The 12 African countries included in this analysis of the causes of TB decline. Drawn with mapchart.net/africa.html.

### HIV and TB epidemic trajectories for 12 countries of southern and eastern Africa

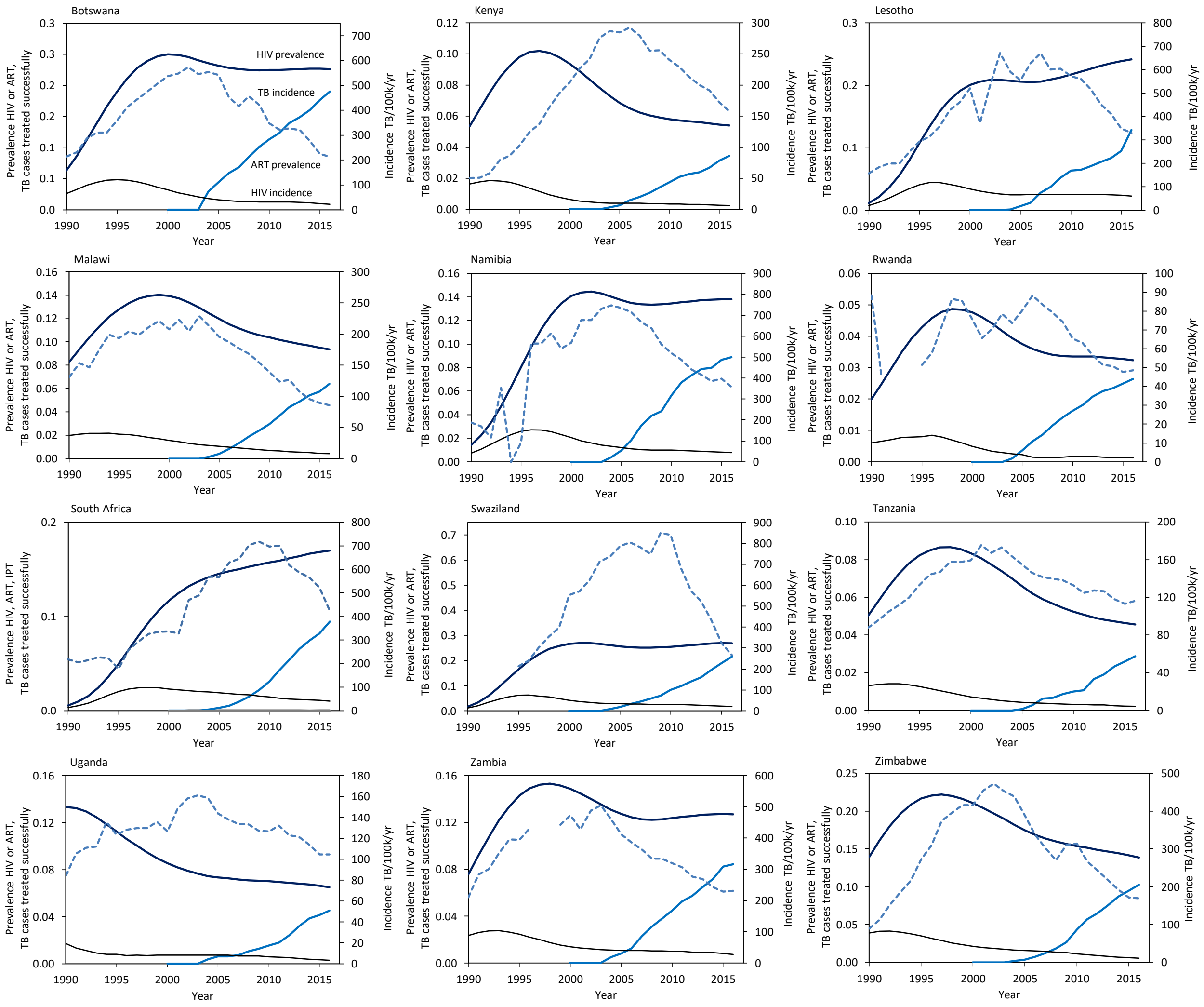


Fig. S2. Time course of the TB and HIV epidemics in 12 countries of southern and eastern Africa, 1990-2016. Continuous lines (y-axis left) show trends in estimated HIV incidence and prevalence, and ART prevalence (coverage in the whole population). Broken lines show the reported TB incidence (y-axis right). Data are from WHO and UNAIDS (*1, 2*).

#### TB case detection and treatment in Mozambique

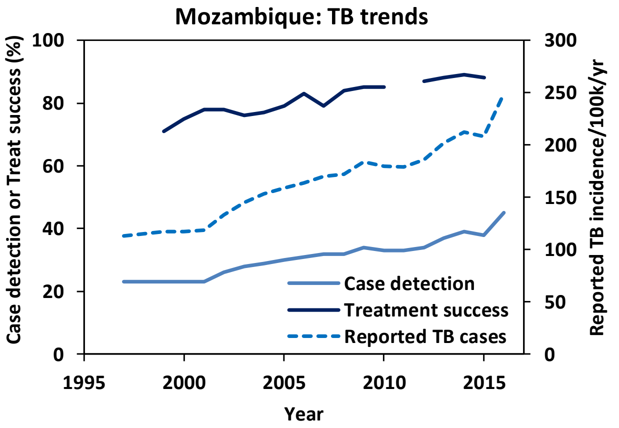


Fig. S3. Reported TB cases, case detection (all TB) and treatment success in Mozambique, 1997-2016

#### Time delay between starting PLHIV on ART and reducing TB incidence

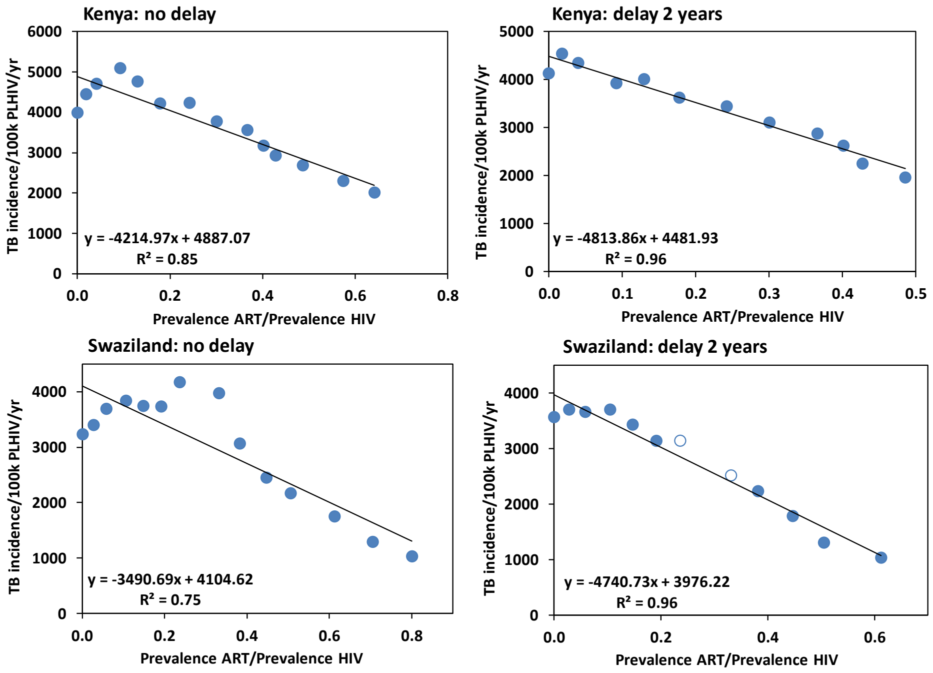


Fig. S4. Fit of the linear regression model without (left) and with (right) a 2-year time delay between TB incidence among PLHIV and the coverage of ART (prevalence of ART/prevalence HIV), illustrated for Kenya and Swaziland. The 2-year delay improves the model fit ($r^2$ on graph) for these and the other ten countries investigated. For Swaziland, the regression model was fitted with interpolated TB incidence data for 2009 and 2010 (open circles, right panel), replacing outlying points for those two years (left panel).

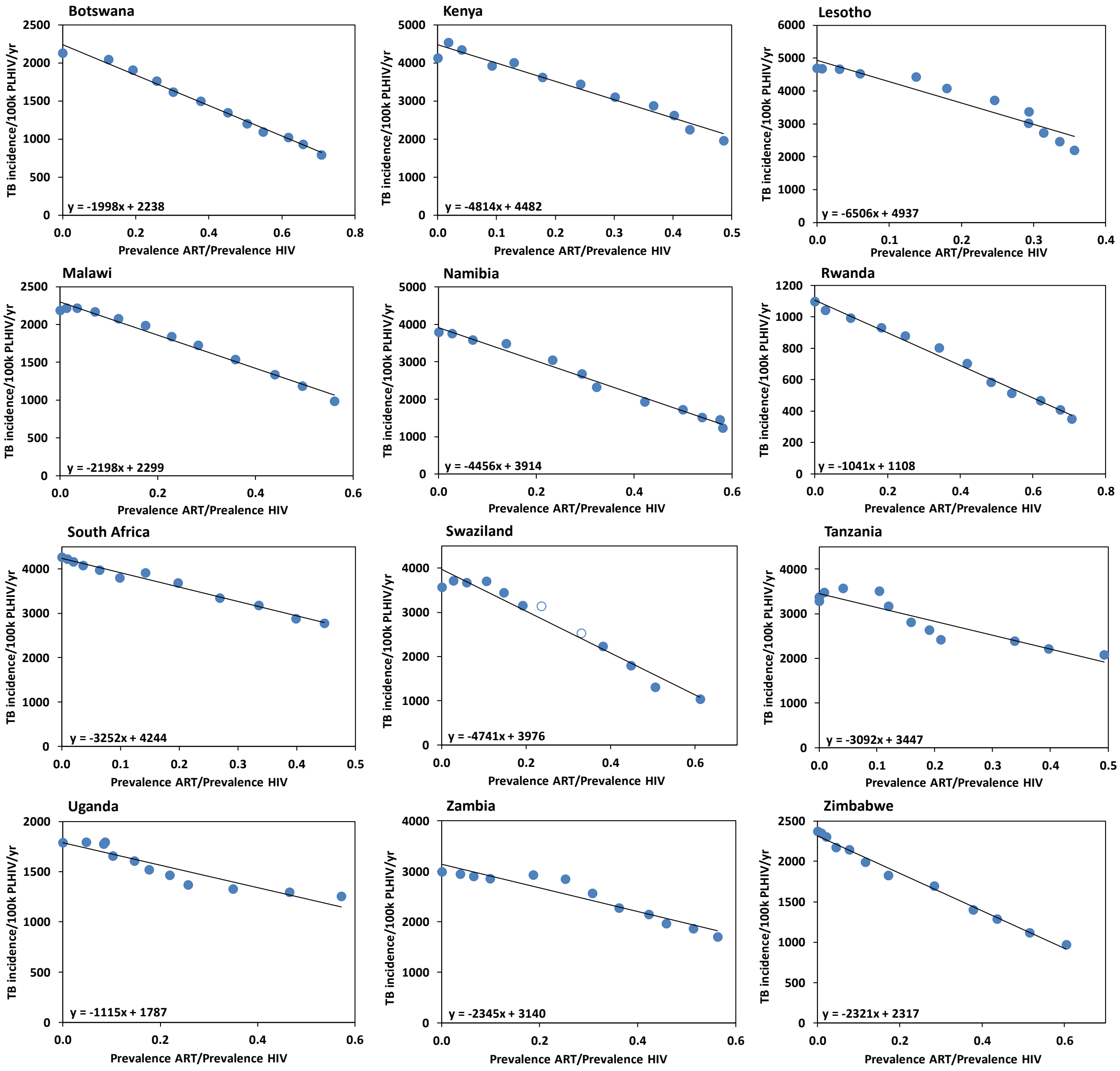


Figure S5. Linear regressions of $T_t/H_{t-\tau}$ on $A$, which give estimates of parameters $\alpha$ and $\beta$ in the model describing the effect of ART on TB incidence (Materials and Methods).

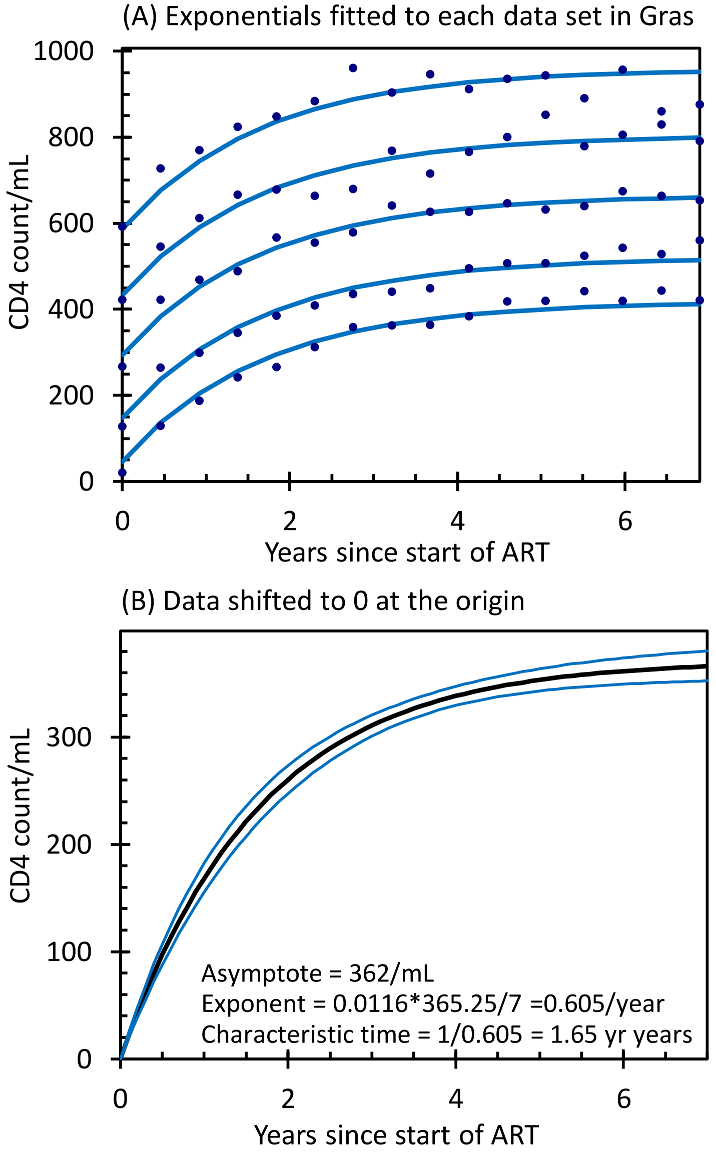


Fig. S6. The characteristic recovery time of CD4 cell count for HIV-infected people started on ART. Data from Gras et al (*3*). (a) The data fitted to exponential curves converging to an asymptote of the form $C_i(t) = a_i(1\text{-}\exp(\varepsilon t))$. $C_i\,(t)$ is the CD4 cell count at time $t$ for people starting at each of 5 CD4 cell count groups at the start of treatment and $\varepsilon$ is the rate of convergence to the asymptote constrained to be the same for all five series of data. (b) The data in (a) but with each data set shifted to an origin of zero and then refitted to show the overall recovery in the CD4 cell count with time with 95% confidence limits on the fitted curve. The recovery in CD4 cell counts after the start of treatment is independent of the value when treatment starts. For the combined data, the recovery rate is $\varepsilon = 0.605$/year; the characteristic time is the inverse, 1.65 years.

#### Regression statistics

Table S1. Regression statistics for the effect of ART on HIV-positive TB (left column) and for the effect of TB detection and treatment on all TB (right column)

Effect of ART on HIV-positive TB

| Botswana | Coefficients | SE | t Stat | P-value | Lower 95% | Upper 95% |
|---|---|---|---|---|---|---|
| α | 2304 | 28.9 | 79.7 | 2.37E-15 | 2239.2 | 2368.0 |
| β | -2121 | 64.2 | -33.0 | 1.53E-11 | -2264.1 | -1977.9 |
| **Kenya** | **Coefficients** | **SE** | **t Stat** | **P-value** | **Lower 95%** | **Upper 95%** |
| α | 4937 | 74.8 | 66.0 | 1.55E-14 | 4770.2 | 5103.5 |
| β | -5857 | 269.8 | -21.7 | 9.61E-10 | -6458.7 | -5256.3 |
| **Lesotho** | **Coefficients** | **SE** | **t Stat** | **P-value** | **Lower 95%** | **Upper 95%** |
| α | 4926 | 123.9 | 39.8 | 2.42E-12 | 4649.7 | 5201.6 |
| β | -6850 | 541.9 | -12.6 | 1.79E-07 | -8057.6 | -5642.9 |
| **Malawi** | **Coefficients** | **SE** | **t Stat** | **P-value** | **Lower 95%** | **Upper 95%** |
| α | 2456 | 18.5 | 132.5 | 1.47E-17 | 2415.2 | 2497.8 |
| β | -2459 | 62.3 | -39.5 | 2.59E-12 | -2598.0 | -2320.6 |
| **Namibia** | **Coefficients** | **SE** | **t Stat** | **P-value** | **Lower 95%** | **Upper 95%** |
| α | 4020 | 57.5 | 69.9 | 8.74E-15 | 3891.4 | 4147.6 |
| β | -4715 | 155.0 | -30.4 | 3.45E-11 | -5060.6 | -4370.0 |
| **Rwanda** | **Coefficients** | **SE** | **t Stat** | **P-value** | **Lower 95%** | **Upper 95%** |
| α | 1197 | 11.9 | 100.2 | 2.39E-16 | 1170.0 | 1223.2 |
| β | -1190 | 27.5 | -43.3 | 1.04E-12 | -1250.9 | -1128.4 |
| **S Africa** | **Coefficients** | **SE** | **t Stat** | **P-value** | **Lower 95%** | **Upper 95%** |
| α | 4084 | 31.2 | 130.8 | 1.68E-17 | 4014.6 | 4153.8 |
| β | -3017 | 137.7 | -21.9 | 8.75E-10 | -3324.1 | -2710.6 |
| **Swaziland** | **Coefficients** | **SE** | **t Stat** | **P-value** | **Lower 95%** | **Upper 95%** |
| α | 4251 | 189.2 | 22.5 | 6.88E-10 | 3829.3 | 4672.7 |
| β | -5277 | 594.9 | -8.9 | 4.71E-06 | -6602.8 | -3951.7 |
| **Tanzania** | **Coefficients** | **SE** | **t Stat** | **P-value** | **Lower 95%** | **Upper 95%** |
| α | 1.4 | 0.1 | 10.0 | 1.62E-06 | 1.1 | 1.7 |
| β | -1.4 | 0.3 | -3.9 | 2.89E-03 | -2.1 | -0.6 |
| **Uganda** | **Coefficients** | **SE** | **t Stat** | **P-value** | **Lower 95%** | **Upper 95%** |
| α | 1827 | 41.9 | 43.6 | 9.75E-13 | 1734.0 | 1921.0 |
| β | -1136 | 156.7 | -7.2 | 2.77E-05 | -1485.0 | -786.5 |
| **Zambia** | **Coefficients** | **SE** | **t Stat** | **P-value** | **Lower 95%** | **Upper 95%** |
| α | 3256 | 57.8 | 56.3 | 7.55E-14 | 3127.2 | 3384.8 |
| β | -2667 | 175.2 | -15.2 | 3.03E-08 | -3057.3 | -2276.8 |
| **Zimbabwe** | **Coefficients** | **SE** | **t Stat** | **P-value** | **Lower 95%** | **Upper 95%** |
| α | 2460 | 34.2 | 71.9 | 6.63E-15 | 2383.4 | 2535.9 |
| β | -2544 | 113.3 | -22.4 | 6.92E-10 | -2796.8 | -2291.7 |

Statistically significant P-vaues are marked in red

Effect of TB detection and treatment on all TB

| | Coefficients | SE | t Stat | P-value | Lower 95% | Upper 95% |
|---|---|---|---|---|---|---|
| r | 0.9 | 0.0 | 29.08 | 5.39E-11 | 0.8 | 1.0 |
| δ | -0.1 | 0.1 | -1.34 | 2.11E-01 | -0.2 | 0.1 |
| **Kenya** | **Coefficients** | **SE** | **t Stat** | **P-value** | **Lower 95%** | **Upper 95%** |
| r | 2.5 | 0.3 | 9.71 | 2.08E-06 | 1.9 | 3.0 |
| δ | -4.1 | 0.7 | -6.12 | 1.13E-04 | -5.6 | -2.6 |
| **Lesotho** | **Coefficients** | **SE** | **t Stat** | **P-value** | **Lower 95%** | **Upper 95%** |
| r | 1.3 | 0.2 | 6.50 | 6.88E-05 | 0.8 | 1.7 |
| δ | -1.1 | 0.6 | -1.81 | 1.00E-01 | -2.4 | 0.2 |
| **Malawi** | **Coefficients** | **SE** | **t Stat** | **P-value** | **Lower 95%** | **Upper 95%** |
| r | 1.2 | 0.2 | 5.45 | 2.81E-04 | 0.7 | 1.6 |
| δ | -0.6 | 0.4 | -1.34 | 2.09E-01 | -1.5 | 0.4 |
| **Namibia** | **Coefficients** | **SE** | **t Stat** | **P-value** | **Lower 95%** | **Upper 95%** |
| r | 1.1 | 0.1 | 11.01 | 6.56E-07 | 0.9 | 1.3 |
| δ | -0.3 | 0.2 | -1.91 | 8.51E-02 | -0.7 | 0.1 |
| **Rwanda** | **Coefficients** | **SE** | **t Stat** | **P-value** | **Lower 95%** | **Upper 95%** |
| r | 1.2 | 0.2 | 6.86 | 4.43E-05 | 0.8 | 1.6 |
| δ | -0.5 | 0.3 | -1.82 | 9.83E-02 | -1.2 | 0.1 |
| **S Africa** | **Coefficients** | **SE** | **t Stat** | **P-value** | **Lower 95%** | **Upper 95%** |
| r | 1.3 | 0.1 | 12.78 | 1.62E-07 | 1.1 | 1.6 |
| δ | -0.7 | 0.2 | -3.27 | 8.38E-03 | -1.2 | -0.2 |
| **Swaziland** | **Coefficients** | **SE** | **t Stat** | **P-value** | **Lower 95%** | **Upper 95%** |
| r | 1.4 | 0.1 | 9.98 | 1.62E-06 | 1.1 | 1.7 |
| δ | -1.4 | 0.3 | -3.91 | 2.89E-03 | -2.1 | -0.6 |
| **Tanzania** | **Coefficients** | **SE** | **t Stat** | **P-value** | **Lower 95%** | **Upper 95%** |
| r | 1.0 | 0.2 | 5.57 | 2.37E-04 | 0.6 | 1.4 |
| δ | -0.3 | 0.6 | -0.46 | 6.55E-01 | -1.7 | 1.1 |
| **Uganda** | **Coefficients** | **SE** | **t Stat** | **P-value** | **Lower 95%** | **Upper 95%** |
| r | 0.9 | 0.1 | 10.98 | 6.72E-07 | 0.7 | 1.1 |
| δ | 0.1 | 0.2 | 0.71 | 4.95E-01 | -0.3 | 0.6 |
| **Zambia** | **Coefficients** | **SE** | **t Stat** | **P-value** | **Lower 95%** | **Upper 95%** |
| r | 0.9 | 0.1 | 13.14 | 1.24E-07 | 0.7 | 1.0 |
| δ | 0.0 | 0.1 | 0.38 | 7.12E-01 | -0.2 | 0.3 |
| **Zimbabwe** | **Coefficients** | **SE** | **t Stat** | **P-value** | **Lower 95%** | **Upper 95%** |
| r | 1.1 | 0.1 | 11.76 | 3.53E-07 | 0.9 | 1.3 |
| δ | -0.5 | 0.2 | -2.59 | 2.68E-02 | -0.9 | -0.1 |
